# Reducing FASN expression sensitizes acute myeloid leukemia cells to differentiation therapy

**DOI:** 10.1101/2020.01.29.924555

**Authors:** Magali Humbert, Kristina Seiler, Severin Mosimann, Vreni Rentsch, Sharon L. McKenna, Mario P. Tschan

**Affiliations:** Institute of Pathology, Division of Experimental Pathology, University of Bern, Bern, Switzerland; TRANSAUTOPHAGY: European network for multidisciplinary research and translation of autophagy knowledge, COST Action CA15138; Graduate School for Cellular and Biomedical Sciences, University of Bern, Bern, Switzerland; Cancer Research, UCC, Western Gateway Building, University College Cork, Cork, Ireland

**Author notes:** Corresponding Author: Magali Humbert, Institute of Pathology, Division of Experimental Pathology, University of Bern, Murtenstrasse 31, CH-3008 Bern, Switzerland.

**Keywords:** FASN, AML, ATRA, TFEB, mTOR, autophagy

## Abstract

Fatty acid synthase (FASN) is the only human lipogenic enzyme available for *de novo* fatty acid synthesis and is often highly expressed in cancer cells. We found that FASN mRNA levels were significantly higher in acute myeloid leukemia (AML) patients than in healthy granulocytes or CD34^+^ hematopoietic progenitors. Accordingly, FASN levels decreased during all-*trans* retinoic acid (ATRA)-mediated granulocytic differentiation of acute promyelocytic leukemia (APL) cells, partially via autophagic degradation. Furthermore, our data suggests that inhibition of FASN expression levels using RNAi or (-)-epigallocatechin-3-gallate (EGCG), accelerates the differentiation of APL cell lines and significantly re-sensitized ATRA refractory non-APL AML cells. FASN reduction promoted translocation of transcription factor EB (TFEB) to the nucleus, paralleled by activation of CLEAR network genes and lysosomal biogenesis. Lysosomal biogenesis was activated, consistent with TFEB transcriptional activation of CLEAR network genes.

Together, our data demonstrate that inhibition of FASN expression in combination with ATRA treatment facilitates granulocytic differentiation of APL cells and may extend differentiation therapy to non-APL AML cells.

## Introduction

While traditional chemotherapy and radiotherapy target highly proliferative cancer cells, differentiation-inducing therapy aims to restore differentiation programs to drive cancer cells into maturation and ultimately into cell death. Differentiation therapies are associated with lower toxicity compared to classical cytotoxic therapies. The success of this therapeutic approach is exemplified by the introduction of all-*trans* retinoic acid (ATRA) in 1985 to treat acute promyelocytic leukemia (APL) (Wang & Chen, 2008). The introduction of ATRA into the treatment regimen changed APL from being one of the most aggressive acute myeloid leukemia (AML) subtypes with a fatal course often within weeks only, to a curable disease with a complete remission rate of up to 95% when combined with anthracycline-based chemotherapy or arsenic trioxide (Wang & Chen, 2008). APL is characterized by translocations involving the C-terminus of the retinoic acid receptor alpha (RARA) on chromosome 17 and genes encoding for aggregate prone proteins. Promyelocytic leukemia (PML)-RARA is the most frequently expressed fusion protein. It is encoded by the translocation t(15;17) and has a dominant negative effect on RARA. RARA transcriptionally regulates multiple biological processes with a key role in differentiation (Germain *et al*, 2006). Several reports suggest a beneficial effect of ATRA in combination therapies in non-APL AML cells (Su *et al*, 2015; Marchwicka *et al*, 2014; Schenk *et al*, 2012). Unfortunately, a variety of intrinsic resistance mechanisms in non-APL AML have been identified such as SCL overexpression, expression of PRAME and epigenetic silencing or mutation of RARA (Rice *et al*, 2004; Bullinger *et al*, 2013; Petrie *et al*, 2009; Altucci & Gronemeyer, 2001). Deciphering the mechanisms active during ATRA-mediated differentiation at the molecular level will support the translation of differentiation therapy to non-APL AML patients. We and others have demonstrated the importance of autophagy in ATRA induced granulocytic differentiation of APL cells (Isakson *et al*, 2010; Wang *et al*, 2011; Jin *et al*, 2018; Humbert *et al*, 2017; Brigger *et al*, 2014b; Orfali *et al*, 2019). Autophagy is an intracellular degradation mechanism that ensures dynamic recycling of various cytoplasmic contents (Feng *et al*, 2014). We thus aim to understand the role of autophagy in granulocytic differentiation and to define key druggable autophagy targets in this process.

Endogenous synthesis of fatty acids is catalyzed by fatty acid synthase (FASN), the only human lipogenic enzyme able to perform *de novo* synthesis of fatty acids (Asturias *et al*, 2005; Maier *et al*, 2006). FASN is frequently overexpressed in a variety of tumor types including leukemias (Pizer *et al*, 1998; Visca *et al*, 2004; Bandyopadhyay *et al*, 2005; Alo *et al*, 1996; Shurbaji *et al*, 1996; Rashid *et al*, 1997; Diaz-Blanco *et al*, 2007) while its expression in healthy adult tissues is low (Weiss *et al*, 1986), with the exception of the cycling endometrium (Pizer *et al*, 1997) and lactating breast (Maningat *et al*, 2009). Interestingly, FASN is upregulated in tumor associated myeloid cells where it activates nuclear receptor peroxisome-proliferator-activated receptor beta/delta (PPARβ/δ) (Park *et al*, 2015), a key metabolic transcription factor in tumorigenesis (Peters & Gonzalez, 2009; Zuo *et al*, 2009). Of note, activation of PPARβ/δ regulates anti-inflammatory phenotypes of myeloid cells in other biological contexts such as atherosclerosis and obesity (Han Jung-Kyu *et al*, 2008; Kang *et al*, 2008; Lee *et al*, 2003; Odegaard *et al*, 2008). We previously reported that (-)-epigallocatechin-3-gallate (EGCG) improved ATRA induced differentiation of APL cells by increasing the expression of death associated protein kinase 2 (DAPK2). Furthermore, EGCG treatment reduce FASN expression levels in selected breast cancer cell lines (Yeh *et al*, 2003). The increased FASN expression in cancer including leukemias, its function in tumor-associated myeloid cells and its link to the differentiation enhancer DAPK2 prompted us to analyze the regulation and function of FASN during myeloid leukemic differentiation.

In the present study, we demonstrate that FASN expression is significantly higher in AML blasts partially due to low autophagic activity in those cells. We show that inhibiting FASN protein expression, but not its enzymatic activity, promotes differentiation of non-APL AML cells. Lastly, we link FASN expression to mTOR activation and inhibition of the key lysosomal biogenesis transcription factor TFEB.

## Material and Methods

### 2.1. Primary cells, cell lines and culture conditions

Fresh leukemic blast cells from untreated AML patients at diagnosis were obtained from the Inselspital Bern (Switzerland) were classified according to the French-American-British classification and cytogenetic analysis. All leukemia samples had blast counts of ~90% after separation of mononuclear cells using a Ficoll gradient (Lymphoprep; Axon Lab AG, Switzerland), as described previously (Tschan *et al*, 2003). Protocols and use of 67 human samples acquired in Bern were approved by the Cantonal Ethical Committee at the Inselspital. The isolation of primary neutrophils (purity 95%) was performed by separating blood cells from healthy donors using polymorphprep (Axon Lab AG, Switzerland). CD34^+^ cells from cord blood or bone marrow were isolated as described (Tschan *et al*, 2003).

The human AML cell lines, HT93, OCI/AML2, MOLM-13 and NB4 were obtained from the Deutsche Sammlung von Mikroorganismen und Zellkulturen GmbH (DSMZ, Braunschweig, Germany). All cell lines were maintained in RPMI-1640 with 10% fetal calf serum (FCS), 50 U/mL penicillin and 50 μg/mL streptomycin in a 5% CO_2_-95% air humidified atmosphere at 37°C. For differentiation experiments, AML cells were treated with 1μM all-*trans* retinoic acid (ATRA; Sigma-Aldrich, Switzerland). Successful granulocyte differentiation was evidenced by CD11b surface expression measured by FACS.

293T cells were maintained in DMEM (Sigma-Aldrich, St. Louis, MO, USA), supplemented with 5% FBS, 1% penicillin/streptomycin, and 1% Hepes (Sigma-Aldrich, Switzerland), and kept in a 7.5%CO_2_-95% air humidified atmosphere at 37°C.

### 2.2 Antibodies

Antibodies used were anti-FASN (3180; Cell Signaling, Switzerland), anti-LC3B (WB: NB600-1384, Novus biological, Switzerland; IF: 3868; Cell Signaling, Switzerland) anti-LAMP1 (14-1079-80; Thermofisher, Switzerland), anti-p62 (HPA003196 ; Sigma-Aldrich, Switzerland), anti-TFEB (4240; Cell Signaling, Switzerland) anti-ULK1(4776; Cell Signaling, Switzerland), anti p-ULK1 (Ser757) (6888; Cell Signaling, Switzerland), anti-ATG13 (6940 ; Cell Signaling, Switzerland), anti-pATG13 (Ser318) (600-401-C49; Rockland, Switzerland), anti p-mTOR (Ser2448) (5536; Cell Signaling, Switzerland), p4E-BP1 (Thr37/46) (2855; Cell Signaling, Switzerland), anti-α-tubulin (3873; Cell Signaling, Switzerland), anti-cleaved PARP (9541; Cell Signaling, Switzerland), anti yH2AX (2577; Cell Signaling, Switzerland) and anti-CD11b-PE (R0841; Dako, Switzerland).

### 2.3 Cell lysate preparation and western blotting

Whole cell extracts were prepared using UREA lysis buffer and 30-60μg of total protein was loaded on a 7.5% or 12% denaturing polyacrylamide self-cast gel (Bio-Rad, Switzerland). Blots were incubated with the primary antibodies in TBS 0.05% Tween-20 / 5% milk overnight at 4°C and subsequently incubated with HRP coupled secondary goat anti-rabbit (7074; Cell Signaling, Switzerland) and goat anti-mouse antibody (7076; Cell Signaling, Switzerland) at 1:5-10,000 for 1 h at room temperature. Blots were imaged using Chemidoc (Bio-Rad, Switzerland) and ImageLab software.

### 2.4 Lentiviral vectors

pLKO.1-puro lentiviral vectors expressing shRNAs targeting *FASN* (sh*FASN*_1: NM_004104.x-1753s1c1 and sh*FASN*_2: NM_004104.x-3120s1c1) were purchased from Sigma-Aldrich. An mCherry-LC3B lentiviral vector was kindly provided by Dr. Maria S. Soengas (CNIO, Molecular Pathology Program, Madrid, Spain). All vectors contain a puromycin antibiotic resistance gene for selection of transduced mammalian cells. Lentivirus production and transduction were done as described (Rizzi *et al*, 2007; Tschan *et al*, 2003). Transduced NB4 cell populations were selected with 1.5 μg/ml puromycin for 4 days and knockdown efficiency was assessed by western blot analysis.

### 2.5 Immunofluorescence microscopy

Cells were prepared as previously described (Brigger *et al*, 2014b). Briefly, cells were fixed and permeabilized with ice-cold 100% methanol for 4 min (LC3B and LAMP1 staining) or 2% paraformaldehyde for 7 min followed by 5 minutes in PBS 0,1% TRITON X-100 (TFEB and tubulin staining) and then washed with PBS. Cells were incubated with primary antibody for 1 h at room temperature followed by washing steps with PBS containing 0.1% Tween (PBS-T). Cells were incubated with the secondary antibody (anti-rabbit, 111-605-003 (Alexa Fluor® 647) 111-096-045 (FITC); anti-mouse, (Cy3) 115-605-003 (Alexa Fluor® 647); Jackson ImmunoResearch, West Grove, PA, USA) for 1 h at room temperature. Prior to mounting in fluorescence mounting medium (S3032; Dako, Switzerland) cells were washed three times with PBS-Tween. Images were acquired on an Olympus FluoView-1000 confocal microscope (Olympus, Volketswil, Switzerland) at 63x magnification.

### 2.6 Acridine Orange staining

Cells were washed 3 times with PBS and resuspended in RPMI 10% FBS containing 5ug/mL Acridine Orange (A3568, Invitrogen, Switzerland) to a concentration of 0.2 x 10^6^ cells per mL. Cells were then incubated at 37°C for 20 min and washed 3 times with PBS. Acridine Orange staining was measured by FACS analysis using a 488nm laser with 530/30 (GREEN) and 695/40 (RED) filters on a FACS LSR-II (BD Biosciences, Switzerland). Data were analyzed with FlowJo software (Ashland, OR, USA). The software derived the RED/GREEN ratio and we compared the distribution of populations using the Overton cumulative histogram subtraction algorithm to provide the percentage of cells more positive than the control.

### 2.7 Nitroblue tetrazolium reduction test

Suspension cells (5 x 10^5^) were resuspended in a 0.2% nitro blue tetrazolium (NBT) solution containing 40ng/ml PMA and incubated 15min at 37°C. Cells were then washed with PBS and subjected to cytospin. Counterstaining was done with 0.5% Safranin O for 5min (HT90432; Sigma Aldrich, Switzerland). The NBT-positive and negative cells were scored under a light microscope (EVOS XL Core, Thermofisher, Switzerland).

### 2.8 Trypan blue exclusion counting

Trypan blue exclusion cell counting was performed to assess cellular growth. 20μL of cell suspension was incubated with an equal volume of 0.4% (w/v) trypan blue solution (Sigma-Aldrich, Switzerland). Cells were counted using a dual-chamber hemocytometer and a light microscope (EVOS XL Core, Thermofisher, Switzerland).

### 2.9 Real-time quantitative RT-PCR (qPCR)

Total RNA was extracted using the RNeasy Mini Kit and the RNase-Free DNase Set according to the manufacturer’s protocol (Sigma-Aldrich, Switzerland). Total RNA was reverse transcribed using all-in-one RT-PCR (BioTool, Switzerland). Taqman® Gene Expression Assays for *BECN1*, *GABARAP*, *STK4,* and *WDR45* were Hs00186838_m1, Hs00925899_g1, Hs00178979_m1 and Hs01079049_g1, respectively. Specific primers and probes for *HMBS* have been already described (Tschan *et al*, 2003). Data represent the mean ± s.d. of at least two independent experiments.

### 2.10 Statistical analysis

Nonparametric Mann-Whitney-U tests were applied to compare the difference between two groups and Spearman Coefficient Correlation using Prism software (GraphPad Software, Inc., Jolla, CA, USA). P-values < 0.05 were considered statistically significant. The error bar on graphs represents the SD of at least two biological replicates performed in two technical replicates

### 3.1. Primary AML blast cells express significantly higher FASN levels compared to mature granulocytes

Cancer cells frequently express high levels of FASN compared to their healthy counterparts (Pizer *et al*, 1998; Visca *et al*, 2004; Bandyopadhyay *et al*, 2005; Alo *et al*, 1996; Shurbaji *et al*, 1996; Rashid *et al*, 1997; Diaz-Blanco *et al*, 2007). We examined *FASN* mRNA expression in an AML patient cohort. *FASN* mRNA levels in AML samples (n=68) were compared to the levels in granulocytes (n=5) and CD34^+^ human hematopoietic progenitor cells (n=3) from healthy donors. We found that *FASN* expression was significantly higher in AML patients compared to healthy granulocytes (p<0.05) (Figure 1A). We obtained similar findings by analyzing *FASN* expression in AML patient data available from the Bloodspot gene expression profile data base (Bagger *et al*, 2016) (Figure 1B). In addition, hematopoietic stem cells from healthy donors express significantly lower *FASN* mRNA transcript levels than AML blasts (Figure 1B). Next, we asked if FASN expression was altered during granulocytic differentiation of APL cells. We analyzed FASN protein expression following ATRA-induced differentiation of two APL cell lines, NB4 and HT93. ATRA treatment resulted in markedly reduced FASN protein levels from day two onwards (Figure 1C). This further suggests that high FASN expression is linked to an immature blast-like phenotype and that ATRA-induced differentiation reduces FASN levels.

**Figure 1:**
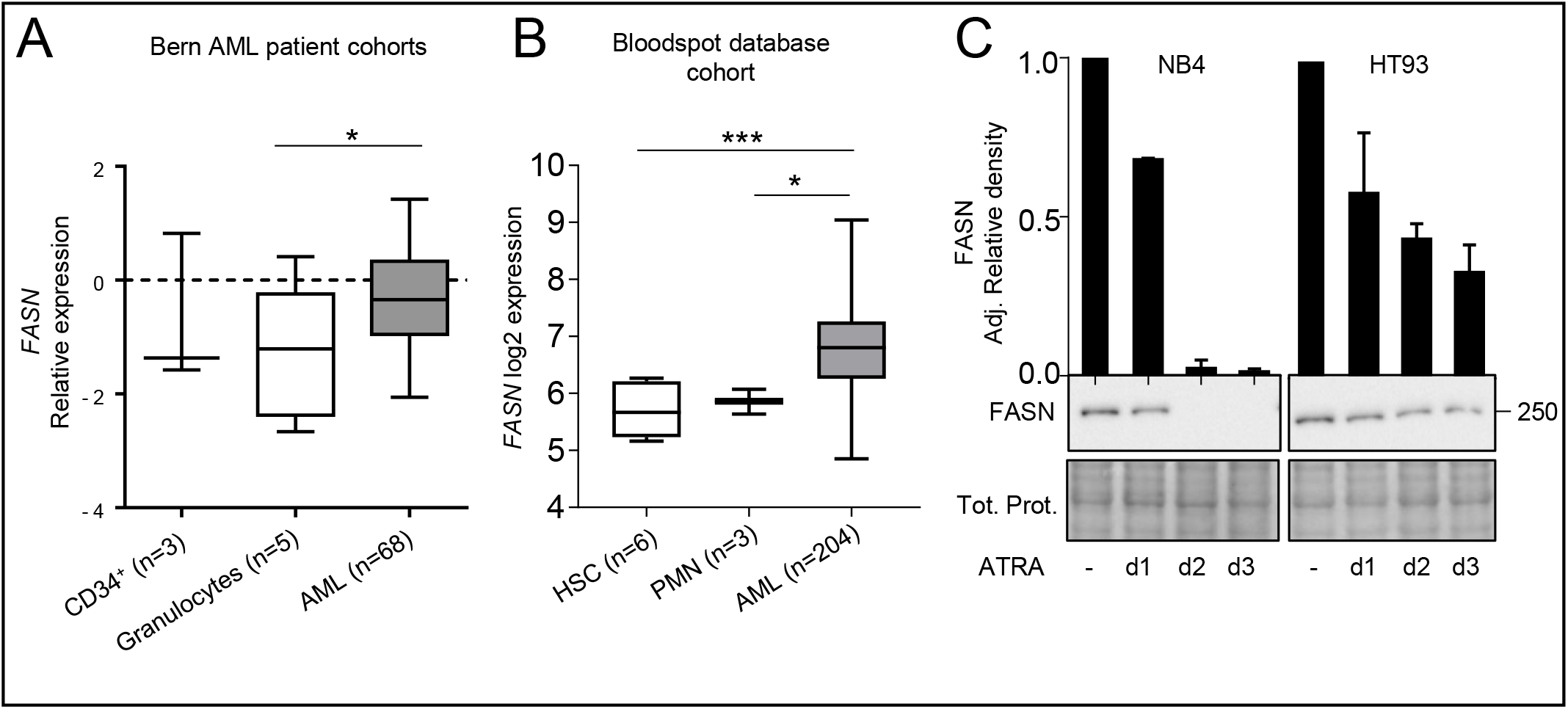
Increased FASN expression is associated with an immature AML blast phenotype. A. FASN mRNA levels in AML blasts, CD34^+^ progenitor cells and granulocytes from healthy donors were quantified by qPCR. All the samples were obtained from the Inselspital, Bern, Switzerland. AML patient cells and granulocytes were isolated using Ficoll gradient density centrifugation. Values are the differences in Ct-values between FASN and the housekeeping gene and ABL1. MNW * p<0.05, ** p<0.0. B. Blood spot data bank analysis of FASN expression in AML blasts compared to granulocytes from healthy donors. MNW * p<0.05, ** p<0.01. C. Western blot analysis of FASN regulation in NB4 and HT93 APL cells upon ATRA treatment at different time points (1, 2 and 3 days). Total protein was extracted and submitted to immunoblotting using anti-FASN antibody. Total protein is shown as loading control. The relative protein expressions were normalized to total protein and quantified using ImageJ software (NIH, Bethesda, MD, USA). Data are represented as a mean (n=3), Error bars: SD.

### 3.2 FASN protein is degraded via macroautophagy during ATRA-induced granulocytic differentiation

We and others have demonstrated that autophagy gene expression is repressed in AML samples compared to granulocytes from healthy donors and that autophagy activity is essential for successful ATRA-induced APL differentiation (Isakson *et al*, 2010; Wang *et al*, 2011; Jin *et al*, 2018; Humbert *et al*, 2017; Brigger *et al*, 2014b; Watson *et al*, 2015; Orfali *et al*, 2015, 2019). The decrease in FASN expression upon ATRA-induced differentiation cannot be explained solely by transcriptional regulation due to the long half-life of this protein (1-3 days)(Volpe & Vagelos, 1976; Weiss *et al*, 1986). Moreover, FASN can be present inside autophagosomes, for instance in yeast and in the breast cancer cell line MCF7 (Dengjel *et al*, 2012; Suzuki *et al*, 2014). Therefore, we hypothesized that ATRA-induced autophagy participates in the degradation of FASN during differentiation of APL cells. To examine whether autophagy is involved in FASN degradation, we treated NB4 cells for 24h with different concentrations of Bafilomycin A1 (BafA1), a specific inhibitor of vacuolar-type H^+^-ATPase (Yamamoto *et al*, 1998; Poole & Ohkuma, 1981), alone or in combination with ATRA. FASN protein was found to accumulate in the presence of BafA1, together with autophagy markers p62 and LC3B-II (Figure 2A). To validate these findings, we utilized NB4 cells stably expressing mCherry-LC3B. Cells were treated with different concentrations of BafA1 with or without ATRA for 24h and FASN as well as LC3B localization was assessed. Endogenous FASN (cyan) showed co-localization with mCherry-LC3B (red) in BafA1 and ATRA treated cells (Figure 2B). In addition, we found colocalization with endogenous FASN (red) and p62 (green) in NB4 parental cells treated with both ATRA and BafA1 for 24h (Figure 2C). It is possible that p62 may help to sequester FASN to the autophagosome. In summary, these data suggest that FASN is a target for autophagic degradation during granulocytic differentiation of APL cells.

**Figure 2:**
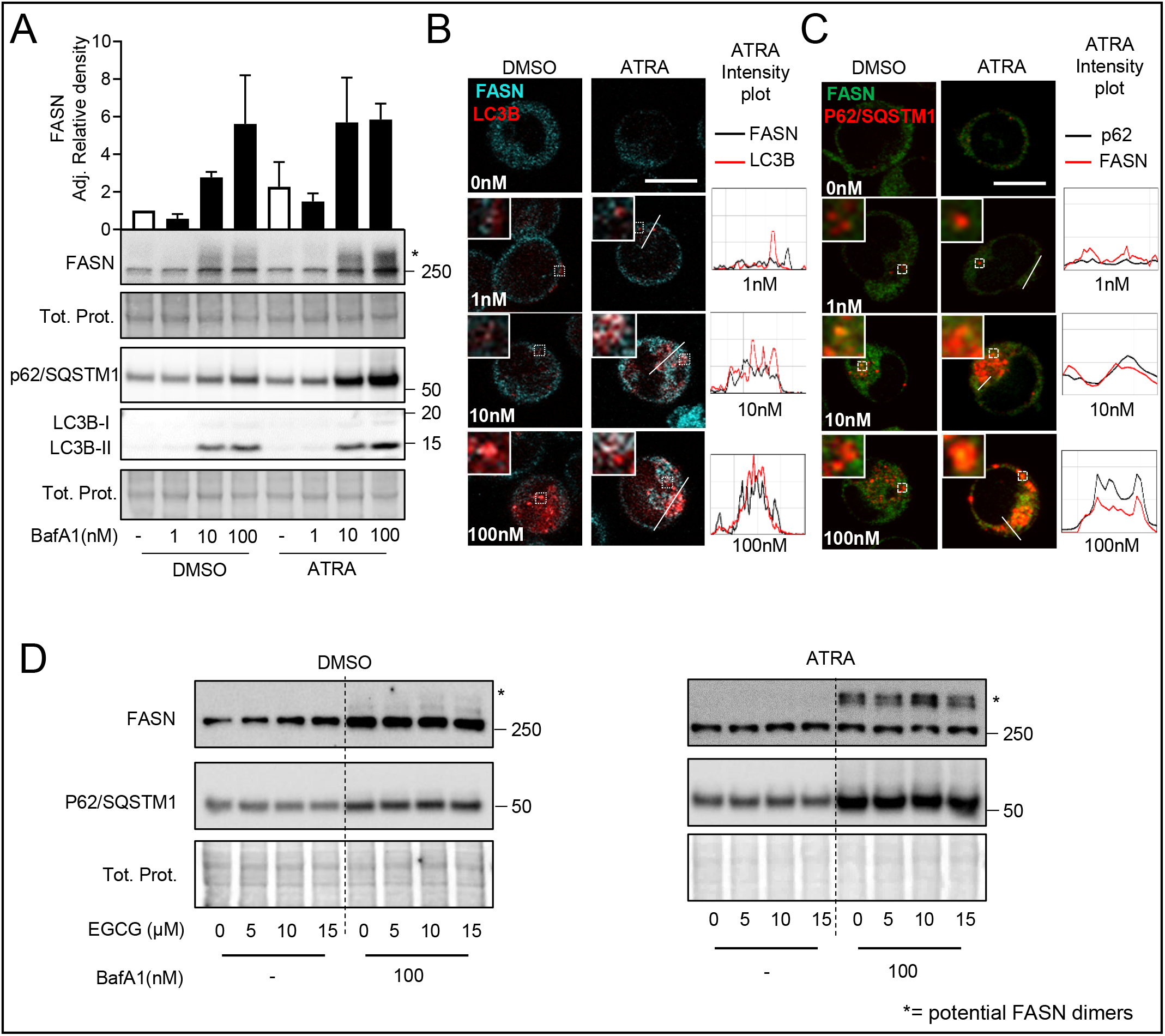
FASN is degraded via autophagy. A. NB4 cells were treated with ATRA and three concentrations of Bafilomycin A1 (BafA1) for 24h. NB4 cells were lysed and subjected to western blot analysis as described in 1C. Quantification of the bands was done using ImageJ software. Data are represented as a mean (n=3), Error bars: SD. B. NB4 cells stably expressing mCherry-LC3B were treated with ATRA and different concentrations of Bafilomycin A1 (BafA1) for 24h. NB4 mCherry-LC3B cells were fixed and stained for endogenous FASN. The colocalization analysis was performed using ImageJ software. Scale: 10μm C. NB4 cells were treated as in 2C and fixed and stained for endogenous FASN and p62. The colocalization analysis was performed using ImageJ software. Scale: 10μm. Results shown are from at least two biological duplicates. (C) EGCG potentiate ATRA-induced FASN degradation by autophagy. (D) FASN and p62 western blot analysis of NB4 cells treated with DMSO (right panel) or ATRA (left panel), in combination with different EGCG (5μM to 15μM) and BafA1 (100nm) concentrations for 24h. Total cell lysates were subjected to western blotting. Quantification of the western blot was done as in Figure 1C.

We have previously shown that EGCG improves the response to ATRA in AML cells by inducing DAPK2 expression, a key kinase in granulocytic differentiation (Britschgi *et al*, 2010). Furthermore, EGCG was reported to decrease FASN expression (Yeh *et al*, 2003) and this was reproducible in our APL cell line model (Supplementary Figure 1A-B). Using different EGCG doses and treatment time points, we confirmed that EGCG improves ATRA induced differentiation in NB4 cells, as evidenced by increased NBT positive cells and CD11b surface expression (Supplementary Figure 1C-E). Importantly, increased differentiation when combining ATRA with EGCG was paralleled by enhanced autophagic activity (Supplementary Figure 1F-G). Autophagy induction was determined by quantifying endogenous, lipidated LC3B-II by western blotting and a dual-tagged mCherry-GFP-LC3B expression construct as described previously (Humbert *et al*, 2017; Gump & Thorburn, 2014; Klionsky *et al*, 2016). Autophagic flux quantification upon EGCG treatment was performed in the presence or absence of BafA1 (Klionsky *et al*, 2016). We found no significant changes in cell death or proliferation measured by DAPI staining and trypan blue exclusion, respectively (Supplementary Figure 2H-I). Co-treating NB4 parental cells with EGCG and ATRA as well as blocking autophagy using BafA1 for 24h, resulted in a higher accumulation of FASN protein (Figure 2D and Supplementary Figure 1J). Interestingly, we saw an accumulation of a band at the molecular weight of FASN dimer (Figure 2D and Supplementary Figure1J). Together, our data demonstrate that FASN can be degraded via autophagy during APL cell differentiation and that co-treatment with EGCG further promotes FASN protein degradation.

### 3.4. Inhibiting FASN protein expression but not its catalytic function accelerates ATRA-induced granulocytic differentiation in APL cell lines

Next, we evaluated the impact of modulating FASN expression and activity on myeloid differentiation. Therefore, we genetically inhibited FASN expression using lentiviral vectors expressing two independent shRNAs targeting *FASN* in the NB4 APL cell line model. Knockdown efficiency was validated by western blotting (Figure 3A). We found that ATRA treatment significantly reduced the doubling time (Supplementary Figure 2A-B) and lowered accumulation of DNA damage as indicated by yH2AX immunofluorescence staining in NB4 *FASN* depleted cells (Supplementary Figure 2C-D). Of note, at steady state conditions, knocking down *FASN* did not affect proliferation compared to control cells. Knocking down *FASN* in NB4 cells resulted in accelerated differentiation into functional granulocytes compared to the control cells as shown by NBT assays (Figure 3B-C) and by CD11b surface expression analysis (Figure 3D). We then assessed the effects of two pharmacological FASN inhibitors, C75 and Orlistat. We used C75 and Orlistat concentrations that do not induce significant cell death (Supplementary Figure 3A-B) or decrease proliferation (Supplementary figure 3C-D) to avoid non-specific effects. Of note, FASN protein levels in APL cells were not reduced by C75 or Orlistat treatment (Supplementary Figure 3E-F1). Unexpectedly, co-treatment of NB4 cells with ATRA and C75 (Figure 3E-G) or Orlistat (Figure 3H-J) did not reproduce the phenotype of the *FASN* knockdown cells. Indeed, cells were differentiating similarly or less compared to control treated cells as demonstrated by NBT assays (Figure 3E-F and Figure 3H-I) and CD11b surface expression (Figure 3G and Figure 3). Therefore, we conclude that the catalytic activity of FASN is not involved in impeding ATRA-mediated differentiation in NB4 cells.

**Figure 3:**
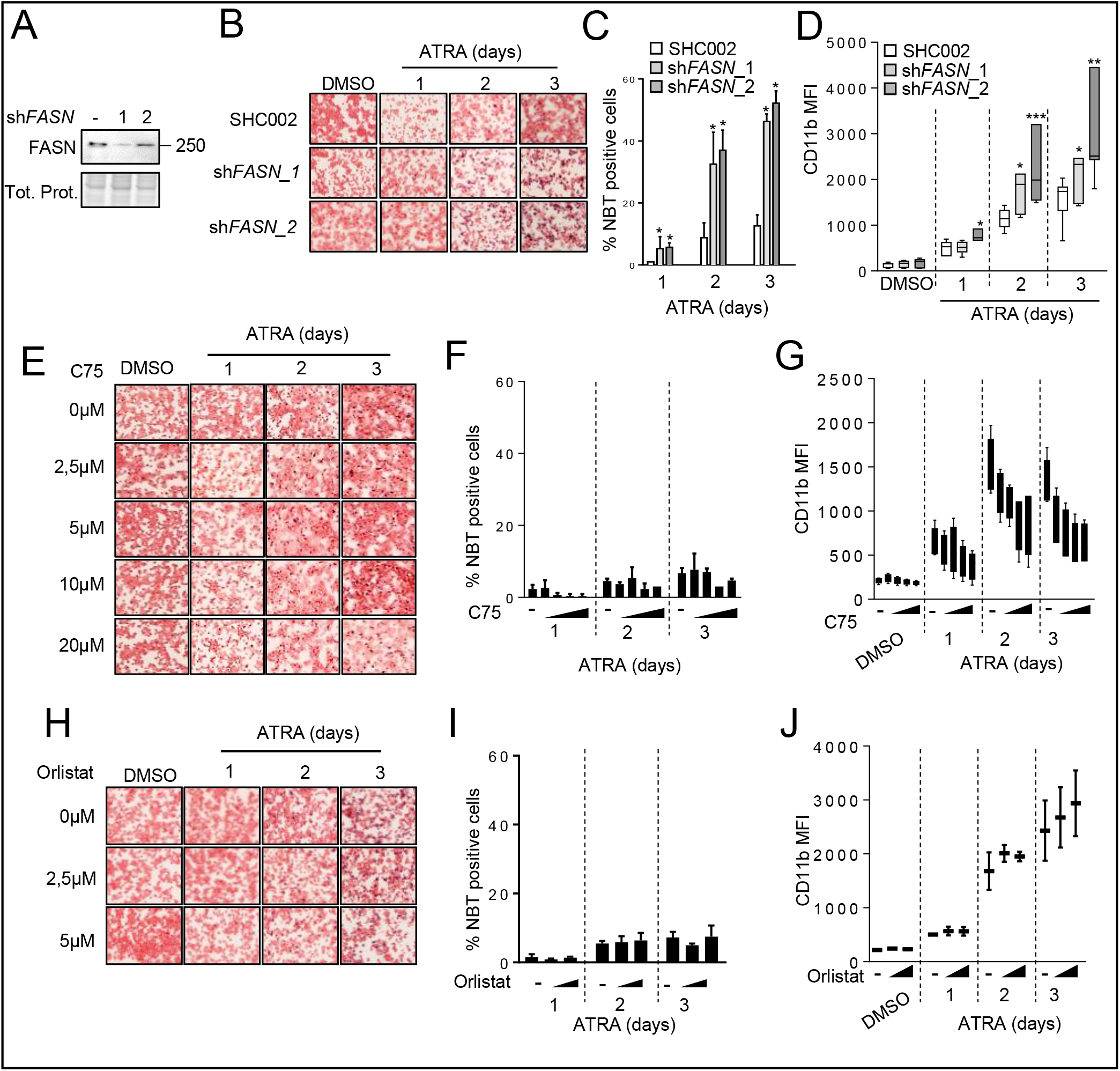
Reducing FASN protein levels improves ATRA-mediated neutrophil differentiation of APL cells. (A-D). NB4 cells were stably transduced with non-targeting shRNA (SHC002) or shRNAs targeting *FASN* (sh*FASN*_1 and sh*FASN*_2) lentiviral vectors and differentiated with 1μM ATRA for 1, 2 or 3 days. A. FASN western blot analysis of control and sh*FASN* (sh*FASN*_1, _2) expressing NB4 cell populations. B-C. NBT reduction in ATRA-treated NB4 control (SHC002) and *FASN* knockdown (sh*FASN*_1, _2) cells. B. Representative images of NBT assays in control and FASN depleted NB4 cells. C. Quantification of the percentage of NBT^+^ cells. D. Flow cytometry analysis of CD11b surface expression NB4 control (SHC002) and *FASN* knockdown (sh*FASN*_1, _2) NB4 cells upon ATRA treatment. E-G. NB4 cells were treated with the indicated C75 concentrations for 3 days in combination with ATRA. E-F. NBT reduction during ATRA-mediated neutrophil differentiation of NB4 control and C75 treated cells. E. Representative images of NBT assays in control and C75 treated NB4 cells upon ATRA-mediated differentiation. F. Quantification of the percentage of NBT^+^ cells. G. Flow cytometry analysis of CD11b surface expression in NB4 control and C75 treated cells upon ATRA-mediated differentiation. H-J. NB4 cells were treated with the indicated Orlistat concentrations for 3 days in combination with ATRA. H-I. NBT reduction during ATRA-mediated neutrophil differentiation of NB4 control and Orlistat treated cells. H. Representative images of NBT assays in control and Orlistat treated NB4 cells upon ATRA-mediated differentiation. I. Quantification of the percentage of NBT^+^ cells. J. Flow cytometry analysis of CD11b surface expression in NB4 control and Orlistat treated cells upon ATRA-mediated differentiation. Data are represented as a mean (n=3), Error bars: SD.

### 3.5. FASN attenuates autophagy by increasing mTOR activity

FASN has been previously reported to promote carcinogenesis by activating mTOR, a master negative regulator of autophagy, via AKT signaling in hepatocellular carcinoma (Hu *et al*, 2016; Calvisi *et al*, 2011). ATRA treatment in APL also reduces mTOR activity leading to autophagy activation (Isakson *et al*, 2010). We therefore hypothesized that FASN may negatively regulate autophagy via mTOR in APL cells, thereby impeding ATRA-induced differentiation. Therefore, we initially confirmed that FASN expression impacts autophagic activity in our system. Autophagy induction was determined by quantifying endogenous LC3B dots formation by immunofluorescence microscopy (IF) after ATRA treatment (Klionsky *et al*, 2016). In order to measure autophagic flux, ATRA treatment was performed in the presence or absence of BafA1 (Figure 4A-B) (Klionsky *et al*, 2016). In addition, we looked at the direct consequences of mTOR activity in ULK1 and ATG13 phosphorylation. ULK1 (ATG1), a key autophagy gene of the initiation complex, is inhibited by mTOR-mediated phosphorylation at Ser757, leading to reduced autophagic activity (Figure 4C) (Kim *et al*, 2011). In line with FASN activating mTOR, lowering *FASN* expression by shRNA (Figure 4D) resulted in decreased mTOR phosphorylation at Ser2448 (Figure 4E) and mTOR-mediated downstream phosphorylation of ULK1 at Ser757 (Figure 4F). Elevated ULK1 activity was confirmed by an increase of ATG13 activating phosphorylation at Ser318 (Figure 4G) (Joo *et al*, 2011; Petherick *et al*, 2015). These results suggest that FASN expression promotes mTOR activity, which in turn enhances autophagy inhibition in AML cells.

**Figure 4:**
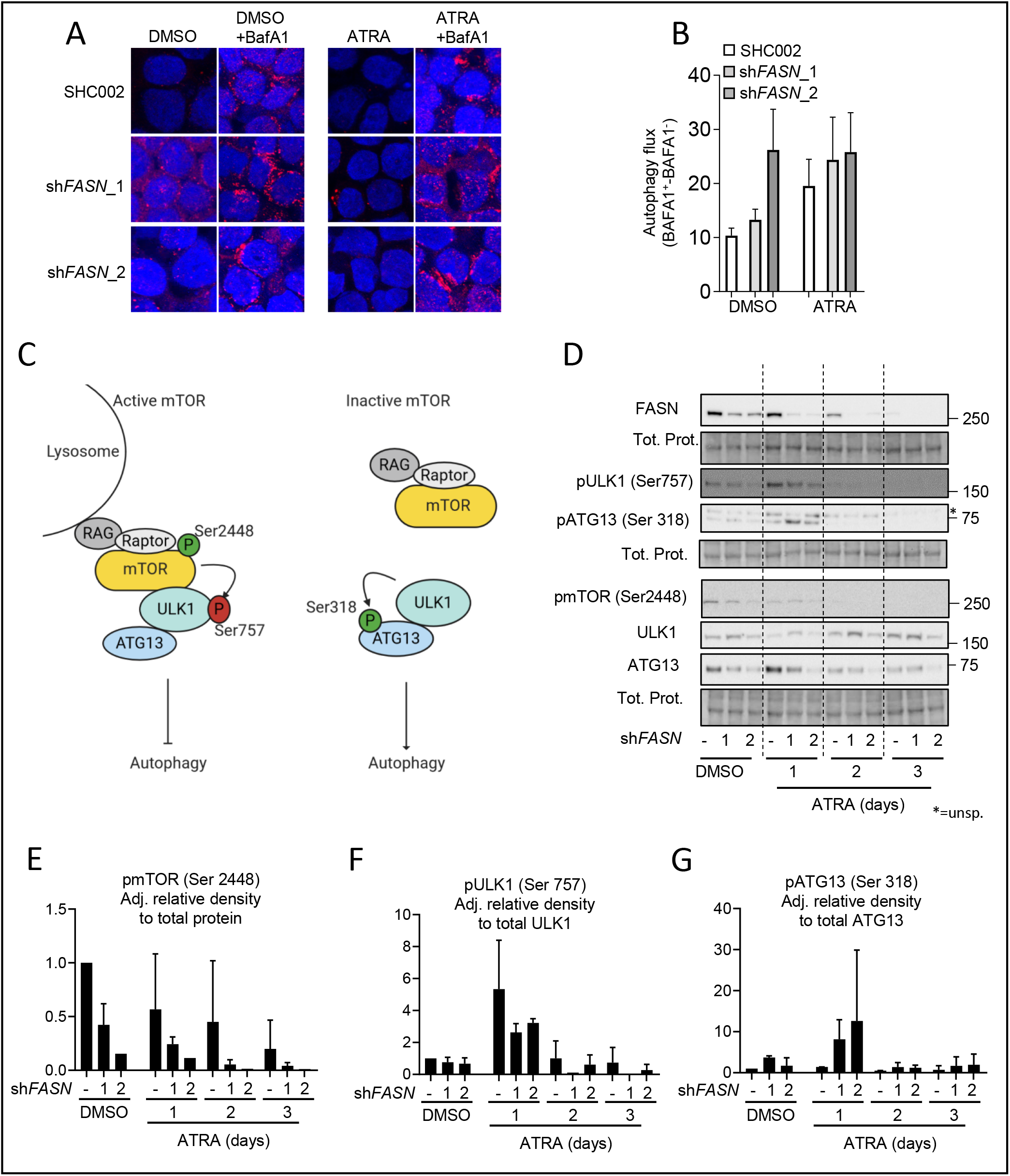
FASN expression is linked to increased mTOR activity. A-C. Autophagy induction in NB4 sh*FASN* cells treated with 1μM ATRA for 24h, in the presence or absence of BafA1 during the last 2h before harvesting. A-B. NB4 control (SHC002) and FASN knockdown (sh*FASN*_1, _2) cells were subjected to LC3B immunofluorescence. A. Representative picture of LC3B punctae in NB4 control (SHC002) and FASN knockdown (sh*FASN*_1, _2) cells. Scale: 10μm B. Quantification of autophagy flux. Three independent experiments were quantified as described in (Humbert *et al*, 2017). C. Scheme of mTOR activity on the ULK1 complex D. NB4 control (SHC002) and FASN knockdown (sh*FASN*_1, _2) cells were treated for 1 to 3 days with ATRA. Total protein was extracted and subjected to immunoblotting using anti-FASN, anti-pmTOR(Ser2448), anti-pULK1(Ser757), anti-ULK1, anti-pATG13(Ser318) and anti-ATG13 antibodies. E-F. Relative protein expressions of two independent experiments were normalized to total protein or the respective non-phosphorylated protein and quantified using ImageJ software (NIH, Bethesda, MD, USA). E. pmTOR(Ser2448) normalized to total protein. F. pULK1(Ser757) normalized to total ULK1. G. pATG13(Ser318) normalized to total ATG13. Results shown are from at least two biological duplicates.

### 3.6. FASN expression negatively affects transcription factor EB (TFEB) activation

mTOR phosphorylates the transcription factor EB (TFEB), a master regulator of lysosome biogenesis, leading to the sequestration of TFEB within the cytoplasm and inhibition of its transcriptional activity (Vega-Rubin-de-Celis *et al*, 2017; Peña-Llopis *et al*, 2011; Roczniak-Ferguson *et al*, 2012; Napolitano *et al*, 2018). TFEB is a key transcriptional regulator of more than 500 genes that comprise the CLEAR (Coordinated Lysosomal Expression and Regulation) network of autophagy and lysosomal genes (Supplementary Figure 4A). A recent study demonstrated the key role of TFEB during ATRA induced differentiation (Orfali *et al*, 2019). We therefore investigated the relationship between FASN and CLEAR network gene expression. Interestingly, the majority of the TFEB downstream targets from the different categories (lysosomal hydrolases and accessory proteins, lysosomal membrane, lysosomal acidification, non-lysosomal proteins involved in lysosomal biogenesis and autophagy) are negatively associated with FASN transcript levels in primary AML patient blasts from TCGA analyzed using the UCSC Xena platform (Goldman *et al*, 2019) and the Blood spot gene expression profiles data base (Bagger *et al*, 2016) (Figure 5A, Supplementary Figures 4B-C Supplementary Table 1-2). Furthermore, analyzing RNA-seq data of NB4 cells treated with ATRA confirmed a reduction of FASN expression paralleled by increased TFEB and TFEB target gene transcript levels (Orfali *et al*, 2019) (Figure 5B). To test if the FASN-mTOR pathway is involved in regulating TFEB activity, we analyzed the cellular localization of TFEB upon ATRA treatment in NB4 control and FASN depleted cells. First, we investigated if TFEB translocates to the nucleus following ATRA treatment and if this translocation is paralleled by an increase in lysosome numbers (LAMP1^+^ dots), assessed by immunofluorescence microscopy (Supplementary Figure 5A-B). Indeed, ATRA treatment resulted in increased LAMP1^+^ dot formation and nuclear translocation of TFEB. Interestingly, TFEB nuclear translocation occurs faster in FASN depleted NB4 cells compared to control cells (Figure 5C), consistent with an increase in LAMP1^+^ dot formation (Figure 5D-E). Furthermore, we treated cells with Acridine Orange to quantify the lysosomal integrity by flow cytometry. Acridine Orange is a cell permeable fluorescent dye that, when excited at 488nm, emits light at 530nm (GREEN) in its monomeric form but shifts its emission to 680nm (RED) when accumulating and precipitating inside lysosomes. Therefore, we measured the RED/GREEN ratio of Acridine Orange stained cells by flow cytometry as previously described (Thomé *et al*, 2016). We found that ATRA treatment shifted the ratio towards the red channel (Supplementary Figure 5C). Reducing FASN expression further accelerated the increase of RED/GREEN ratio indicating enhanced lysosome biogenesis (Figure 5G-H). These results suggest that FASN expression impairs TFEB translocation to the nucleus and therefore reduces lysosome biogenesis

**Figure 5:**
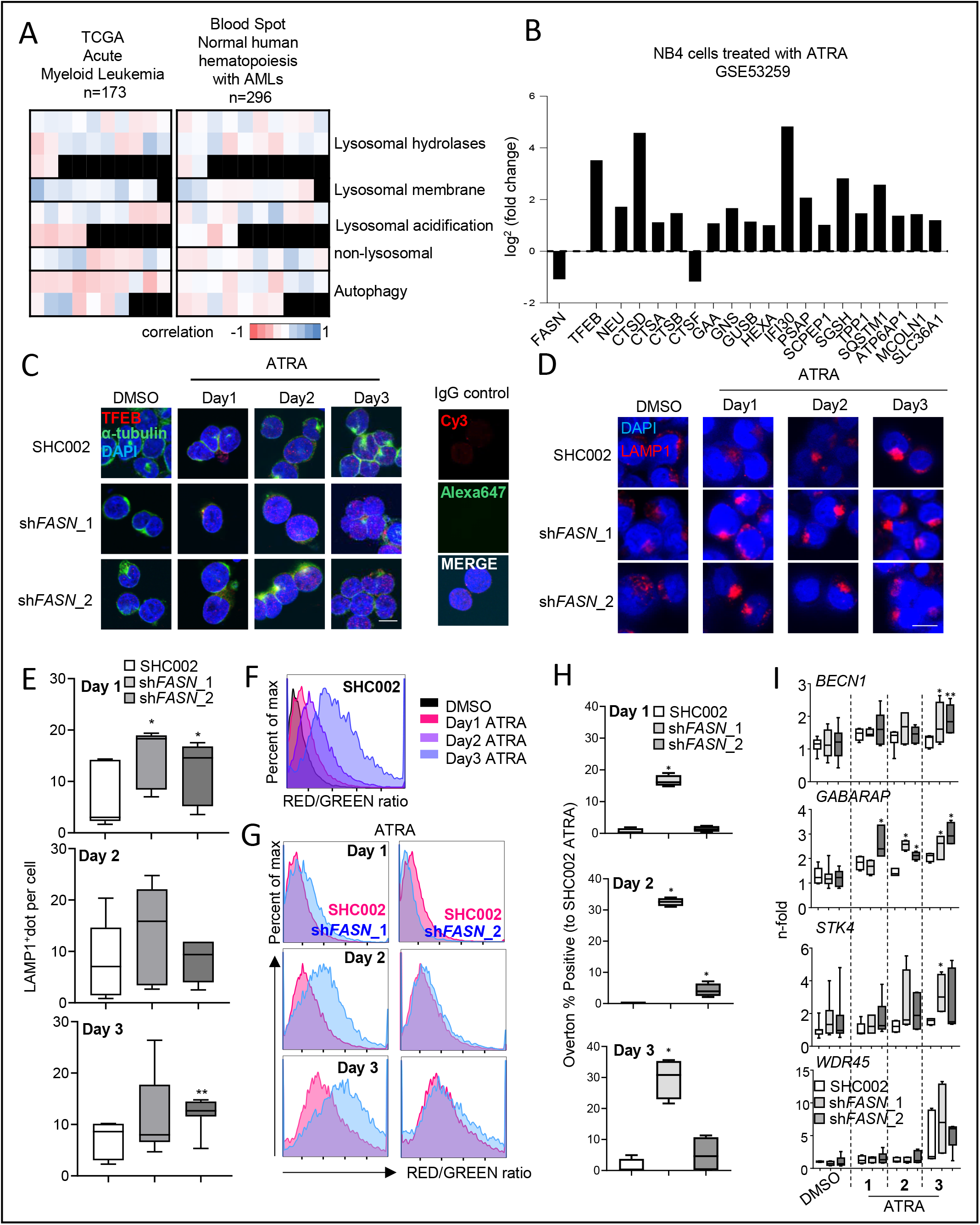
FASN expression negatively associates with TFEB activity. A. Heatmaps of the correlation between FASN and TFEB target genes extracted from the TCGA-AML cohort analyzed by the UCSD xena platform and from the bloodspot data bank (Spearman, p values in Supplementary Table1 and 2). B. mRNA sequencing data of NB4 cells treated with ATRA. Relative expression of *FASN*, *TFEB* and TFEB transcriptional targets involved in lysosomal function and biogenesis are shown. C-H. NB4 control (SHC002) and *FASN* knockdown (sh*FASN*_1, _2) cells were treated for 1 to 3 days with ATRA. (C) Immunofluorescence microscopy of endogenous TFEB (red) and α-tubulin (green). IgG staining was used as negative control. Nuclei were stained with DAPI (blue). (D) Immunofluorescence microscopy of endogenous LAMP1 (red). Nuclei were stained with DAPI (blue). Scale: 10μm (E) LAMP1 punctae quantification of cells shown in D. Scale: 10μm (F-H) Acridine Orange staining. (F) Histogram representation of the ratio between RED and GREEN of NB4 control (SHC002) cells treated as described in 6C. (G) Representative histogram of NB4 control (SHC002) and *FASN* knockdown (sh*FASN*_1, _2) cells treated as in 6C. (H) Overton percentage positive quantification of the RED/GREEN ratio of NB4 control (SHC002) and *FASN* knockdown (sh*FASN*_1, _2) cells treated with ATRA at indicated times. (I) Evaluation of *BECN1*, *GABARAP*, *STK4* and *WDR45* mRNA transcripts was done by qPCR. Values were normalized to the HMBS housekeeping gene. Results shown are from at least two biological duplicates.

We then evaluated the effect of FASN expression on the transcription of the following TFEB target genes: *BECN1*, *GABARAP*, *STK4* and *WDR45*. All 4 TFEB targets showed increased expression upon ATRA treatment, in line with previous studies (Orfali *et al*, 2015; Brigger *et al*, 2013, 2014a) (Figure 5I). Knock down of *FASN* led to a further increase in the expression of 3/4 TFEB targets analyzed (Figure 5I). These results suggest that FASN retardation of TFEB translocation to the nucleus attenuates CLEAR network gene transcription.

Then, we tested whether we can obtain similar results by lowering FASN protein levels using EGCG. Using different EGCG concentrations, we found a decrease in mTOR phosphorylation at Ser2448 (Figure 6A), an increase of TFEB translocation to the nucleus (Figure 6B), an increase of LAMP1^+^ vesicles (Supplementary Figure 6A-B) and an increase of the RED/GREEN ratio in Acridine Orange stained cells similar to the results seen in FASN-depleted APL cells (Supplementary Figure 6C-E). In addition, we found an upregulation of 3/4 TFEB target genes in presence of EGCG in line with our FASN knockdown experiments (Figure 6C).

**Figure 6:**
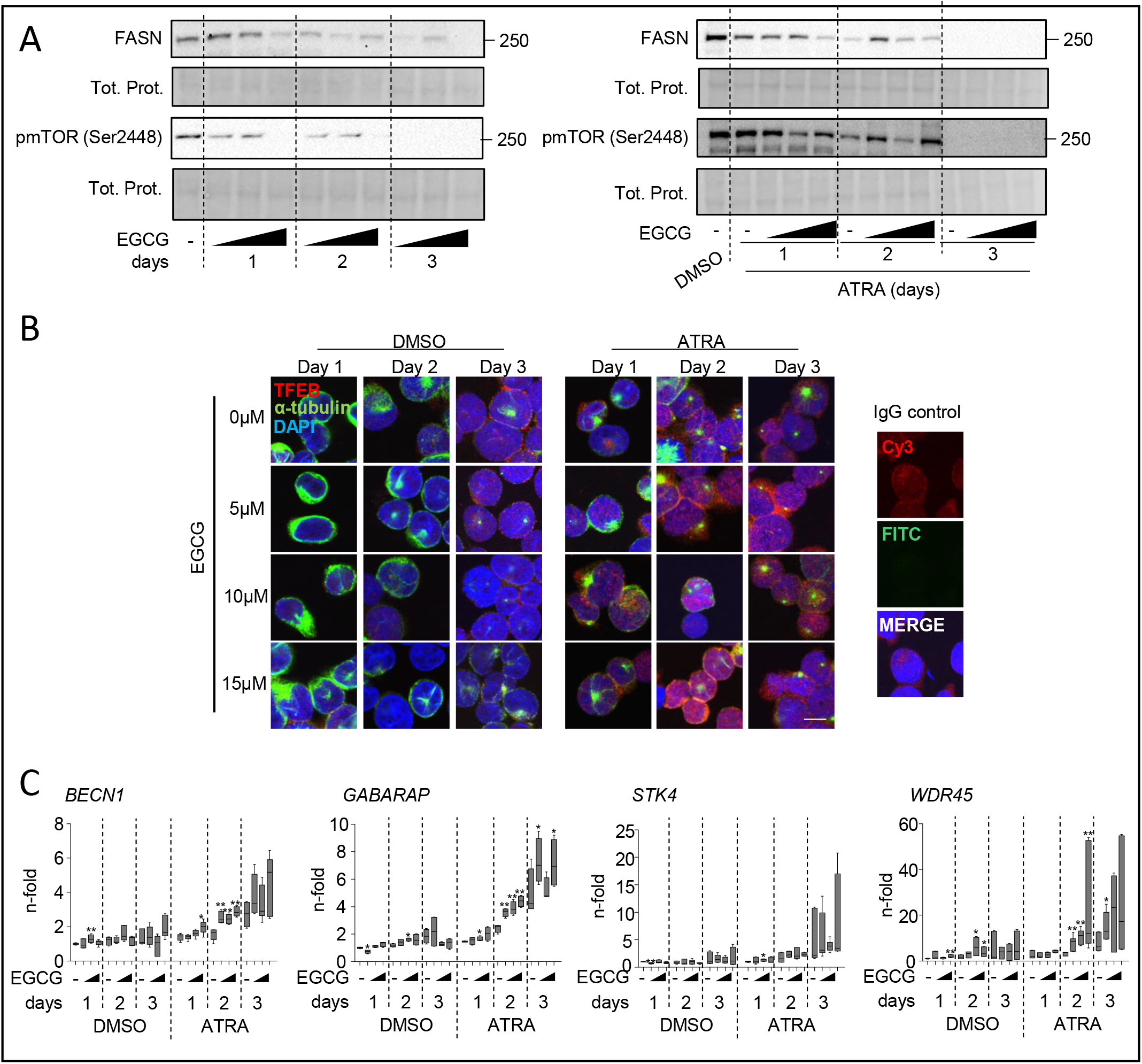
EGCG treatment accelerates TFEB translocation to the nucleus and improves lysosome biogenesis. NB4 and HT93 APL cells were treated for 1 to 3 days with ATRA in combination with indicated concentrations of EGCG. Cells were then subjected to (A) Western blot analysis of FASN and pmTOR(Ser2448), (B) TFEB endogenous immunofluorescence. B. Representative pictures of TFEB in NB4 cells treated with ATRA and different concentrations of EGCG (5μM, 10μM and 15μM), Scale: 10μm. (C) Evaluation of *BECN1*, *GABARAP*, *STK4* and *WDR45* mRNA transcripts was done by qPCR. Values were normalized to the HMBS housekeeping gene. Results shown are from at least two biological duplicates.

Together, these data suggest that high FASN expression results in lower autophagic activity and decreased lysosomal capacity due to increased mTOR activity causing TFEB inhibition.

### 3.7. Lowering FASN expression improves ATRA therapy in non-APL AML cell lines by inhibiting the mTOR pathway

Given the fact that APL cells treated with EGCG demonstrated improved response to ATRA therapy, we asked if EGCG can be beneficial to other AML subtypes that are refractory to ATRA treatment. We and others previously demonstrated a positive impact of co-treating HL60 AML cells, a non-APL AML cell line that responds to ATRA, with EGCG and ATRA (Britschgi *et al*, 2010; Moradzadeh *et al*, 2018; Lung *et al*, 2002). Therefore, we tested if ATRA-refractory AML cell lines with different genetic backgrounds, namely MOLM-13 (FLT3-ITD^+^) and OCI/AML2 (DNMT3A R635W mutation), would respond to ATRA in combination with EGCG. Both cell lines showed increased granulocytic differentiation upon the combination treatment as shown by CD11b surface expression (Figure 7A-B). In addition, MOLM-13 and OCI/AML2 showed an increase of RED/GREEN ratio when stained with Acridine Orange (Figure 7C-D). Furthermore, co-treatment with ATRA and EGCG led to a decrease in mTOR activity as seen by a decrease in mTOR (Ser2448) and ULK1 (Ser757) phosphorylation. In MOLM-13, it was paralleled by an increase of ATG13 (Ser318) phosphorylation (Figure 7E-F). We further confirmed these data by genetically inhibiting *FASN* in MOLM-13 (Figure 8A) and OCI/AML2 (Figure 8B) cells. Depleting FASN in both cell lines caused an increase of CD11b surface expression after 3 days of ATRA treatment (Figure 8C-D), coupled with an increased RED/GREEN ratio when stained for Acridine Orange (Figure 8E-F) and lower mTOR activity (Figure 8G-H). Interestingly, we found more variation in lysosomal compartment changes between the experimental duplicates upon ATRA when cells were treated with EGCG (Figure 7D and 7F) than in the knockdown cells (Figures 8F and 8H), perhaps reflecting the lower specificity of EGCG.

**Figure 7:**
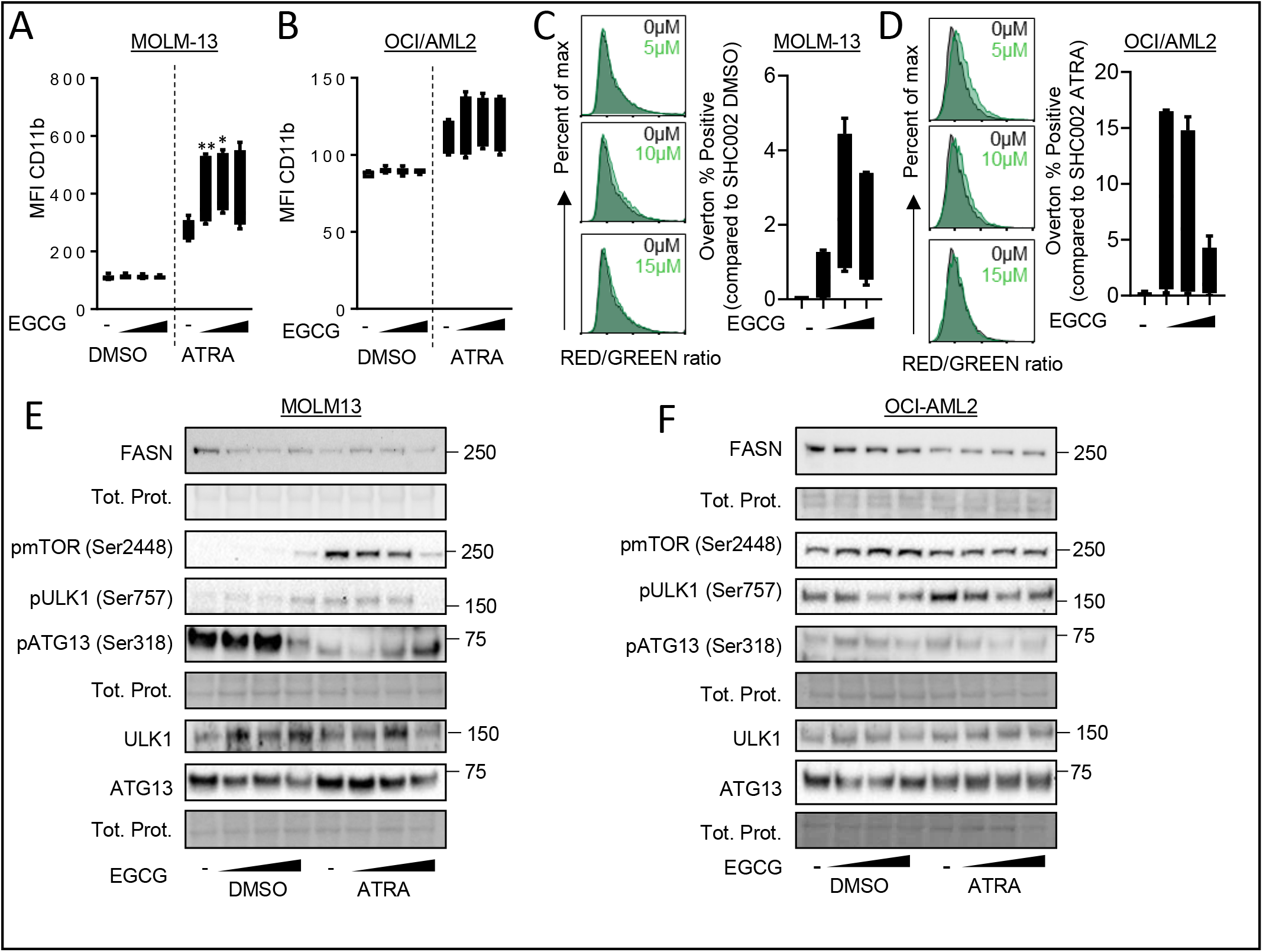
Lowering FASN protein expression levels improves ATRA therapy in non-APL AML cells. A-F: MOLM-13 (A, C, E), and OCI-AML2 (B, D, F) cells were treated with ATRA and different concentrations of EGCG (5μM, 10μM and 15μM) for 3 days (n=3). (A-B) CD11b surface staining was analyzed by flow cytometry. Box blot represent the median fluorescence intensity (MFI) of CD11b positive cells. C-D. Acridine Orange staining. Analysis was performed as in Figure 6E-G. E-F. Total protein extracted from MOLM-13 and OCI-AML2 cells treated as in 7A/B were subjected to immunoblotting using anti-FASN, anti-pmTOR(Ser2448), anti-pULK1(Ser757), anti-ULK1, anti-pATG13(Ser318) and anti-ATG13 antibodies. Results shown are from at least two biological duplicates.

**Figure 8:**
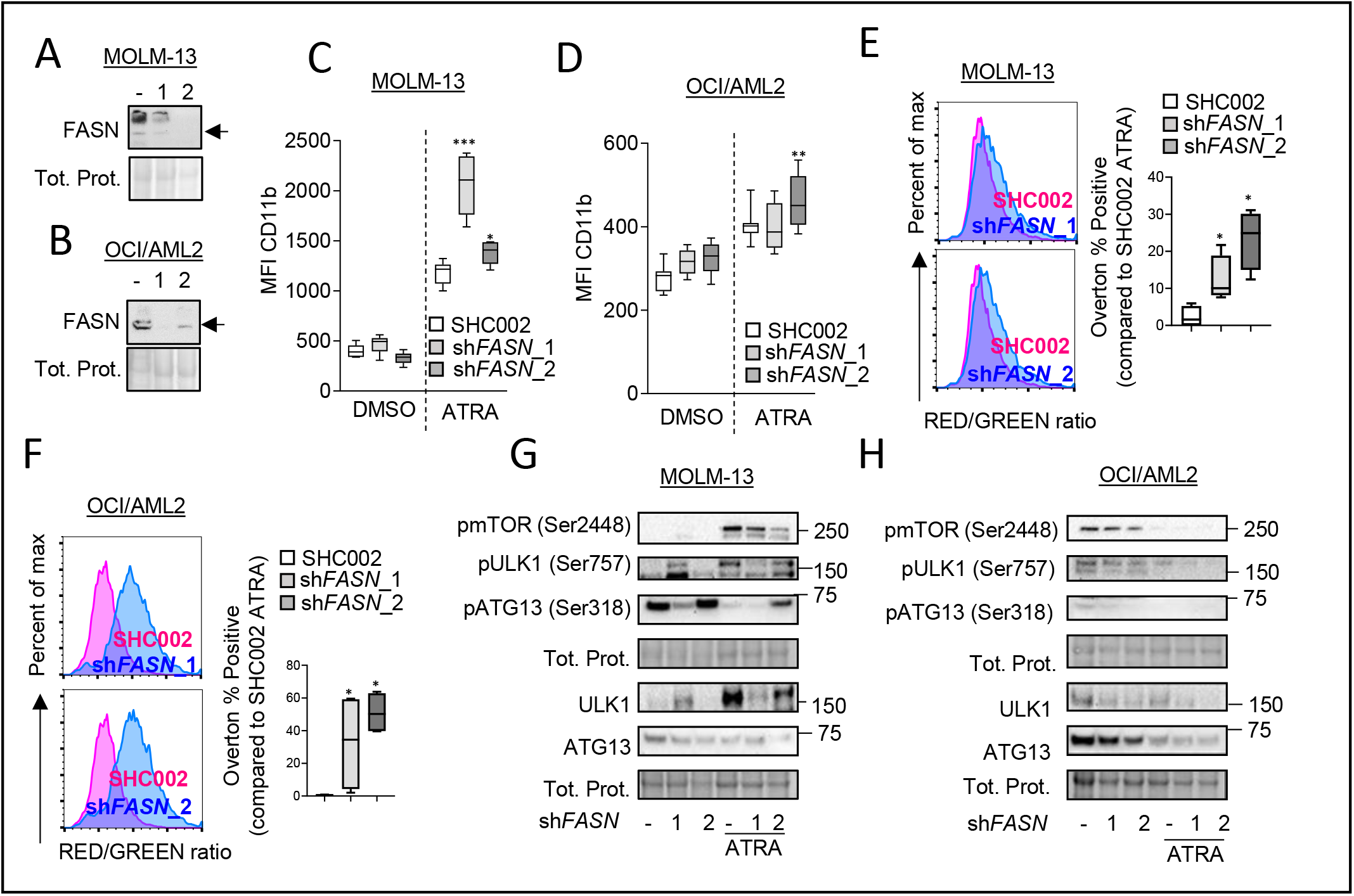
MOLM-13 and OCI/AML2 were stably transduced with 2 independent shRNA targeting *FASN* (n=3) A-B.: *FASN* knockdown efficiency was validated in MOLM-13 (A) and OCI/AML2 (B) by western blotting. C-H. MOLM-13 and OCI/AML2 control and FASN knockdown cells were treated with ATRA for 3 days. (C-D) CD11b surface marker expression was analyzed as in A-B. (E-F) Acridine Orange staining analysis was performed as in 7E-G. (G-H) Western blot analysis of total protein was extracted and subjected to immunoblotting using anti-FASN, anti-pmTOR(Ser2448), anti-pULK1(Ser757), anti-ULK1, anti-pATG13(Ser318) and anti-ATG13 antibodies. Results shown are from at least two biological duplicates.

Together, our data suggest that reducing FASN expression can increase lysosomal biogenesis and improve the differentiation of non-APL AML cells.

## Discussion

In this study, we aimed at further dissecting the function of fatty acid synthase in AML cells and, in particular, its potential role in the differentiation of immature AML blasts. We showed that knocking down *FASN* accelerated ATRA-induced differentiation, while inhibition of its enzymatic function by pharmacological inhibitors such as C75 or Orlistat had no effect. Furthermore, we found that FASN expression activates mTOR resulting in sequestration of TFEB to the cytoplasm. Importantly, inhibiting *FASN* expression, in combination with ATRA treatment, improved differentiation therapy in non-APL AML cells.

Several studies demonstrated a tumor suppressor role of autophagy in AML cells. Autophagy can support degradation of leukemic oncogenes in AML such as FLT3-ITD and PML-RARA (Larrue *et al*, 2016; Rudat *et al*, 2018; Isakson *et al*, 2010). Furthermore, activation of mTORC1 is crucial for leukemia cell proliferation at least partially due to its inhibitory effect on autophagy (Hoshii *et al*, 2012; Watson *et al*, 2015). Accordingly, inhibition of autophagy leads to acceleration of MLL-ENL AML leukemia progression *in vivo* (Watson *et al*, 2015). Our results indicate that increased FASN expression might be a key activator of mTORC1 in AML. Surprisingly, we found that reducing FASN protein levels, but not inhibition of catalytic function, promotes ATRA-induced differentiation. Recently, Bueno *et al.* demonstrated that FASN is key during the transformation from 2-to 3-dimensional growth of cancer cells. This transformation step does not depend on the FASN biosynthetic products palmitate, further hinting to important non-catalytic functions of FASN in carcinogenesis (Bueno *et al*, 2019). Further studies on the interplay between FASN, mTOR and autophagy in AML transformation, progression and therapy resistance are warranted to improve our understanding of cell fate decisions and could potentially open new avenues to tackle this disease with improved differentiation therapies.

We further confirmed that EGCG positively impacts on cellular differentiation in additional AML subtypes *in vitro* (Britschgi *et al*, 2010; Moradzadeh *et al*, 2018; Lung *et al*, 2002). Searching for potential mediators of the positive effects of EGCG observed during ATRA-induced differentiation, we previously found that EGCG induces expression of the Ca^2+^/calmodulin-regulated serine/threonine kinase DAPK2. DAPK2 plays a major role in granulocytic differentiation and decreased DAPK2 expression in APL cells can be restored by ATRA and EGCG treatment (Rizzi *et al*, 2007; Britschgi *et al*, 2010; Humbert *et al*, 2017). DAPK2 also negatively regulates mTOR via phosphorylation of raptor at Ser721 as shown in HeLa cells (Ber *et al*, 2015). Therefore, a potential impact of FASN on DAPK2 activity in a leukemic context warrants further investigation.

Interestingly, treating APL cells with ATRA had a negative effect on FASN protein levels (Figure 1C), and we demonstrated that ATRA-induced autophagy contributes to FASN protein degradation. Furthermore, FASN reduction led to increased lysosomal biogenesis suggesting a negative feedback loop between autophagy and FASN. It is reasonable to hypothesize that the more AML cells differentiate the more they become competent to degrade long-lived proteins including FASN. In addition, inhibiting mTOR using Rapamycin or Everolimus accelerates differentiation of APL cells (Jin *et al*, 2018; Isakson *et al*, 2010). While PI3K/AKT/mTOR pathways are activated in about 80% of AML cases, mTOR inhibitors had only modest effects in AML therapy (Mirabilii *et al*, 2018; Tabe *et al*, 2017). Furthermore, despite its role in leukemia cells, mTOR activity is crucial for hematopoietic stem cell (HSC) proliferation and self-renewal potential (Ghosh & Kapur, 2016). Therefore, targeting FASN with low expression in healthy progenitor cells would allow activation of autophagy in AML cells sparing healthy HSC cells in the bone marrow. In our hands, EGCG treatment only demonstrated partial effects regarding improved differentiation when compared to knocking down FASN in non-APL AML cells. Therefore, a more specific FASN expression inhibitor is needed to improve differentiation therapy in non-APL AML patients.

Indeed, it would be of interest to study the transcriptional regulation of FASN to influence its expression in autophagy deficient cells. Consistently, there are several studies showing that *FASN* transcription is positively affected by retinoic acids (Roder *et al*, 1996; Roder & Schweizer, 2007). However, transcription induction is not mediated by a classic retinoic acid responsive element (RARE) but rather by indirect influence of retinoic acid via *cis*-regulatory elements. Since this involves different cofactors it is tempting to speculate that transcriptional activation might switch to repression depending on the cellular context including specific retinoid-binding proteins and cofactors. We previously found that members of the KLF transcription factor family are often deregulated in primary AML patient samples. Among the different KLF family members downregulated in AML, particularly *KLF5* turned out to be essential for granulocytic differentiation (Humbert *et al*, 2011; Diakiw *et al*, 2012; Li *et al*, 2019). KLF5 forms a transcriptionally active complex with RAR/RXR heterodimers (Lv *et al*, 2013; Kada *et al*, 2008). Interestingly, ectopic expression of KLF5 in U937 non-APL AML cell line was sufficient to significantly increase ATRA-induced differentiation (Shahrin *et al*, 2016). We hypothesize that KLF5 negatively regulates *FASN* transcription in AML cells via the RAR/RXR complex.

In summary, our data suggest that inducing FASN protein degradation is likely to be beneficial for differentiation therapy of non-APL AML cells as this will impede mTOR and promote TFEB transcriptional activity and autophagy. Furthermore, high FASN expression in AML is partially based on attenuated autophagy activity in this disease.

## Acknowledgments

Deborah Shan-Krauer is gratefully acknowledged for excellent technical support. We thank Dr. MS Soengas for providing a mCherry-LC3B lentiviral vector. This study was supported by grants from Swiss Cancer Research (KFS-3409-02-2014 to MPT, MD-PhD 03/17 Scholarship to KS), the Swiss National Science Foundation (31003A_173219 to MPT), the Berne University Research Foundation (45/2018, to MPT), the University of Bern initiator grant, the Bernese Cancer League, “Stiftung für klinisch-experimentelle Tumorforschung”, and the Werner and Hedy Berger-Janser Foundation for Cancer Research (to MH). SMCK is supported by Breakthrough Cancer Research.

## Author Contribution

MH, KS, SM, and VR performed the experimental research. MH, KS, SMCK and MPT drafted the article. MH designed the project. MPT gave final approval of the submitted manuscript.

