## Supplementary Figure Legends for "Reducing FASN expression sensitizes acute myeloid leukemia cells to differentiation therapy"

**Supplementary Figure 1:** EGCG treatment reduces FASN expression and increases autophagy activity upon ATRA treatment. NB4 cells were treated with different EGCG concentration (5µM, 10µM or 15µM) up to 3 days together with ATRA. A-B. Total lysate from ATRA- (A) and ATRA+ (B) treated cells were subjected to immunoblotting using anti-FASN. Total protein is shown as a loading control. The relative protein expressions were normalized to total protein and quantified using ImageJ software (NIH, Bethesda, MD, USA). C-D. NBT reduction in ATRA-induced NB4 control and C75 treated cells C. most representative images of NBT assay. D. Quantification of the percentage of $NBT^+$ cells. E. Flow cytometry analysis of CD11b surface expression NB4 control and EGCG treated cells upon ATRA. F. NB4 cells were treated with EGCG at indicated concentrations in presence or absence of ATRA (1µM) and BafA1. Total lysate was subjected to immunoblotting using anti-p62 and anti-LC3B. Total protein is showed as a loading control. G. NB4 cells were stably transfected with a mCherry-EGFP-LC3B construct and treated with different EGCG concentration together with ATRA for the indicated time point. Ratio between mCherry and EGFP was measured by flow cytometry. H-I. NB4 parental cells were treated as in A and B. H. non fixed cells were stained with DAPI as a cell death marker. I. NB4 cells were counted using typan blue exclusion method at day 0, 1, 2, 3 and 6. J. FASN and p62 western blot analysis of NB4 cells treated with DMSO or ATRA, in combination with different EGCG (5µM to 15µM) and BafA1 (100nm) concentrations for 24h. Total cell lysates were subjected to western blotting. Biological duplicate of Figure 2 D.

**Supplementary Figure 2:** NB4 sh*FASN* cells were treated for 3 days with ATRA. A-B cells were counted using typan blue exclusion method at day 0, 1, 2, 3 and 6. A.

growth curve extracted from the counting, B. doubling time of the NB4 sh*FASN* cells in control conditions or ATRA. C-D. γH2AX foci was analyzed by immunofluorescence upon ATRA treatment at day 1, 2 and 3. C. Representative pictures. Scale: 10µm. D. Quantification of foci per cells.

**Supplementary Figure 3:** NB4 parental cells were treated with either C75 at concentration 2.5µM, 5µM, 10µM or 20µM (A, C, E) or Orlistat 2.5µM or 5µM (B, D, F). A-B non fixed cells were stained with DAPI as a cell death marker. C-D. NB4 cells were counted using typan blue exclusion method at day 0, 1, 2, 3 and 6. E-F. total cell lysate was subjected to immunoblotting using anti-FASN. Total protein is showed as a loading control.

**Supplementary Figure 4:** A. Schematic of TFEB activity. (B-C) Analysis of gene expression of *FASN*, *TFEB* and the CLEAR network in (B) TCGA-AML data set and (C) Blood Spot normal human hematopoiesis with AMLs.

**Supplementary Figure 5:** NB4 parental cells were treated for 3 days with 1µM ATRA and cells were then subjected to (A) TFEB endogenous immunofluorescence (B) LAMP1 endogenous immunofluorescence and (C) Acridine Orange staining. A. representative picture of TFEB in NB4 cells. Scale: 10µm. B. representative picture of LAMP1 in NB4. Scale: 10µm. C. Histogram representation or the ratio between RED and GREEN of NB4 cells treated with ATRA.

**Supplementary Figure 7:** (A-B) LAMP1 endogenous immunofluorescence, and (C-E) Acridine Orange staining. A. Representative pictures of LAMP1 in NB4 cells treated with ATRA and different concentrations of EGCG. Scale: 10µm. B. LAMP1 punctae quantification in ATRA and EGCG treated cells at indicated time points. C. Histogram representation or the ratio between RED and GREEN of NB4 cells treated with EGCG at indicated time and concentration. D. Representative histogram of NB4 cells treated with EGCG and ATRA at indicated time points and concentrations. E. Overton percentage positive quantification of the RED/GREEN ratio of NB4 cells treated with DMSO (upper panel) or ATRA (lower panel) at indicated times and in combination with indicated EGCG concentrations.

**Supplementary Table 1:** Spearman Correlation coefficient of TCGA data set between FASN expression and CLEAR network

**Supplementary Table 2:** Spearman Correlation coefficient of the bloodspot data set between FASN expression and CLEAR network
