## Supplementary Figures for "Reducing FASN expression sensitizes acute myeloid leukemia cells to differentiation therapy"

### Supplementary fig 1

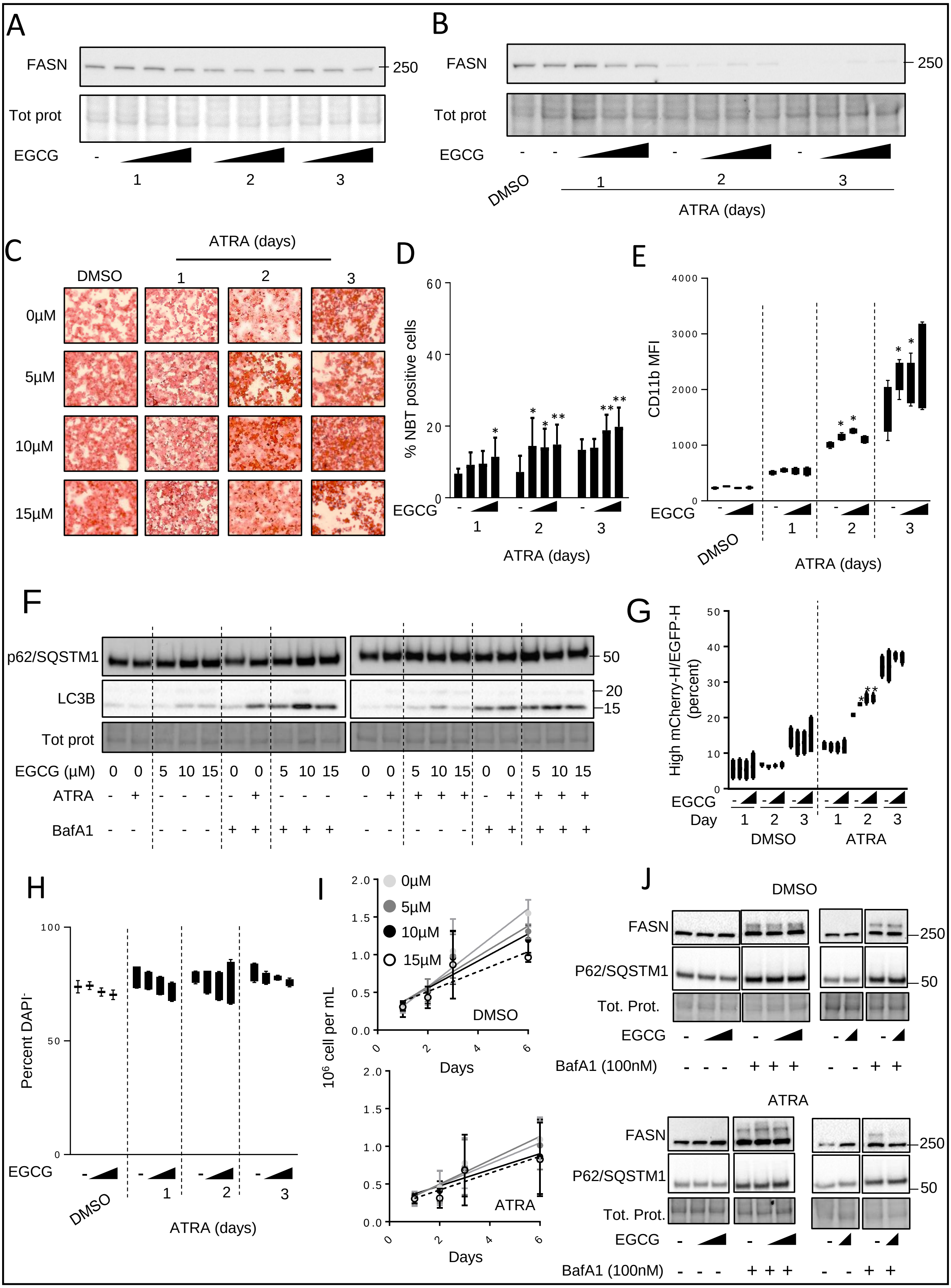

### Supplementary fig 2

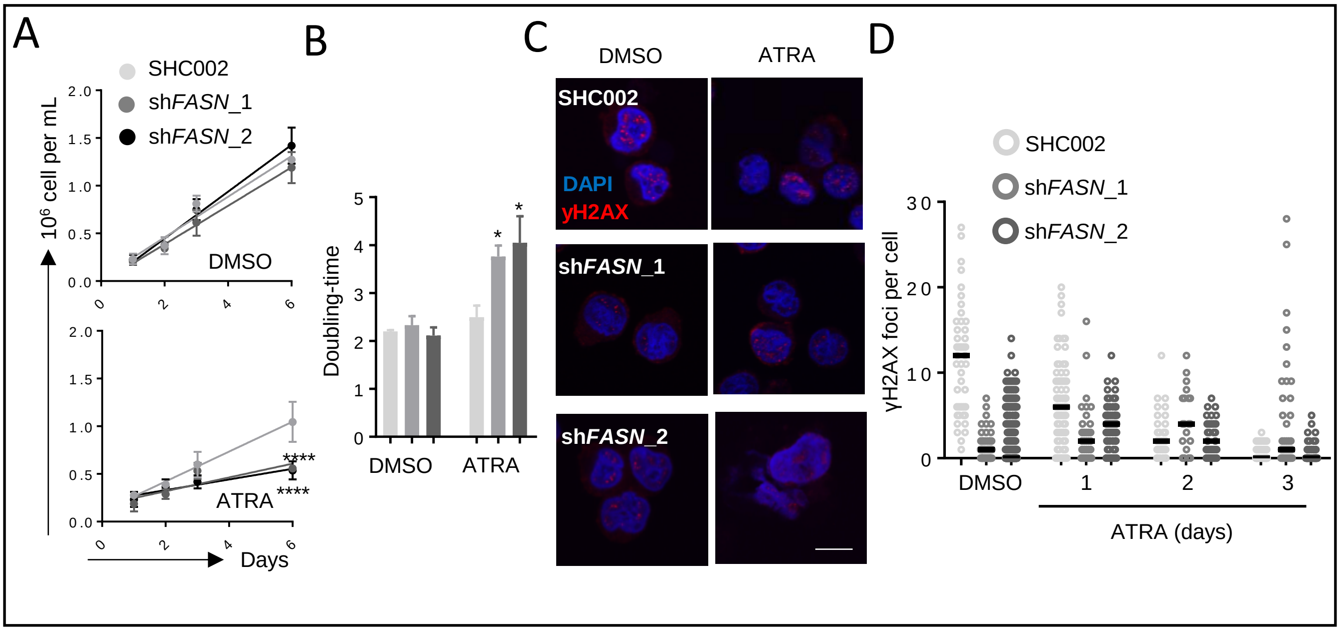

### Supplementary fig 3

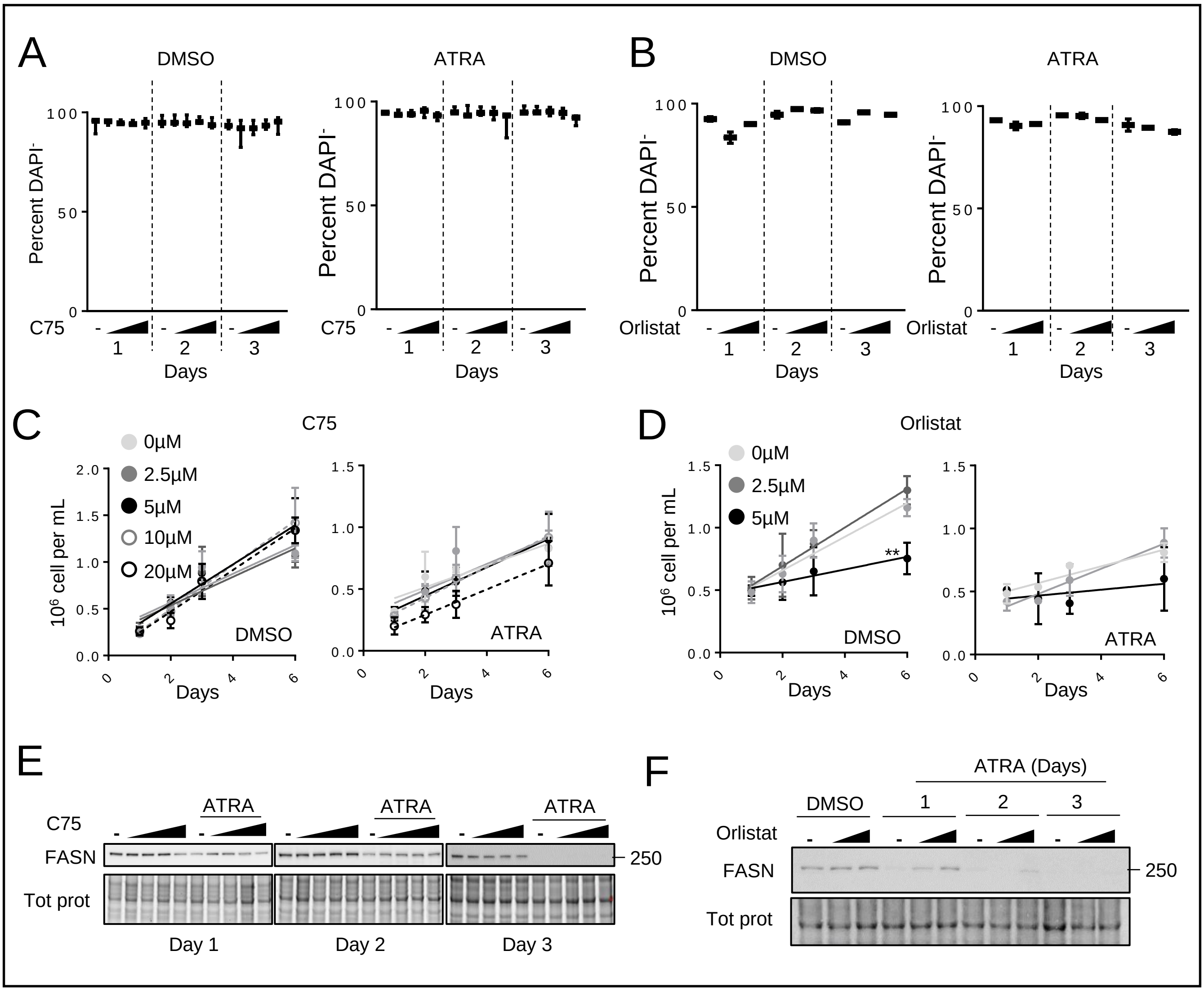

### Supplementary fig 4

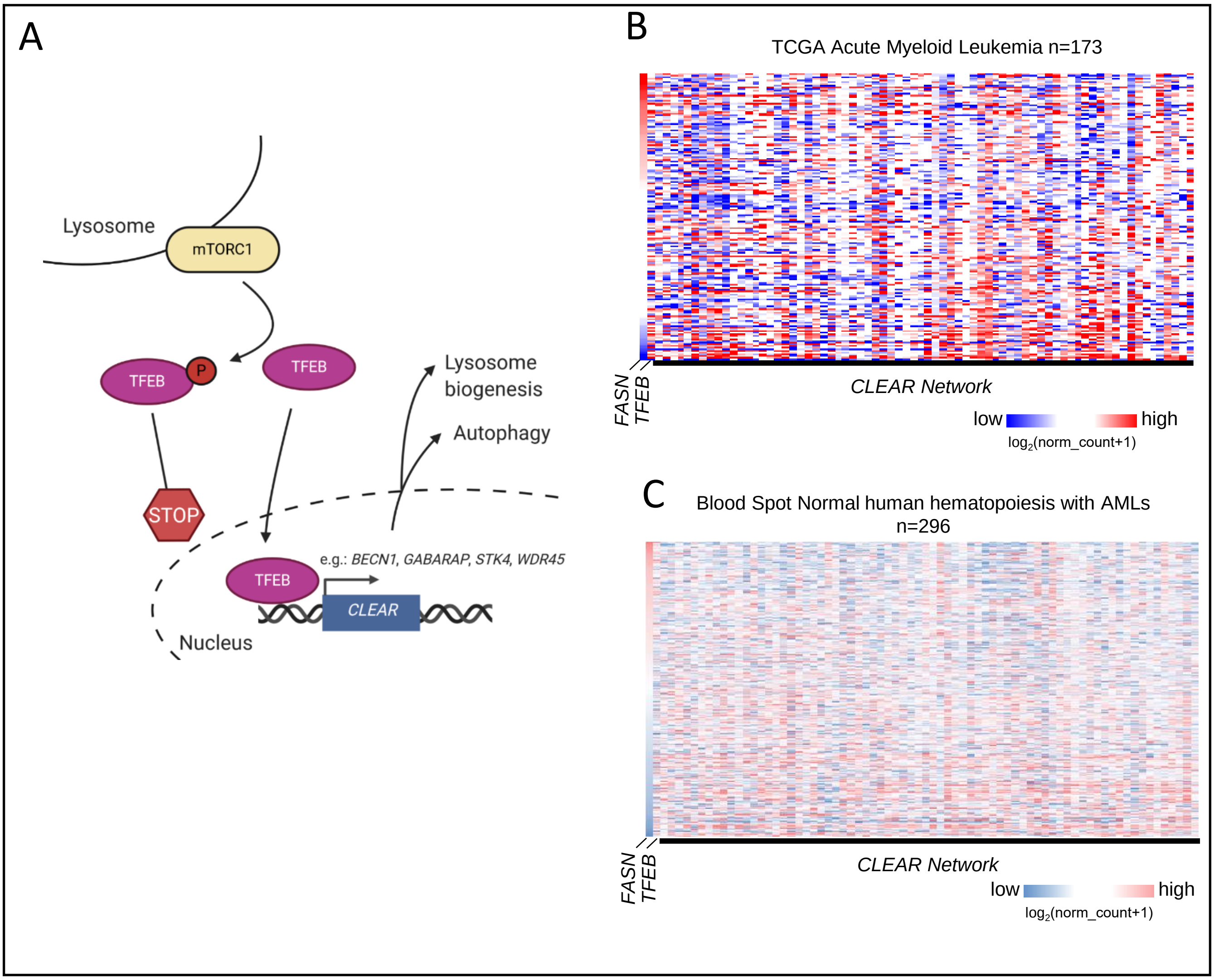

### Supplementary fig 5

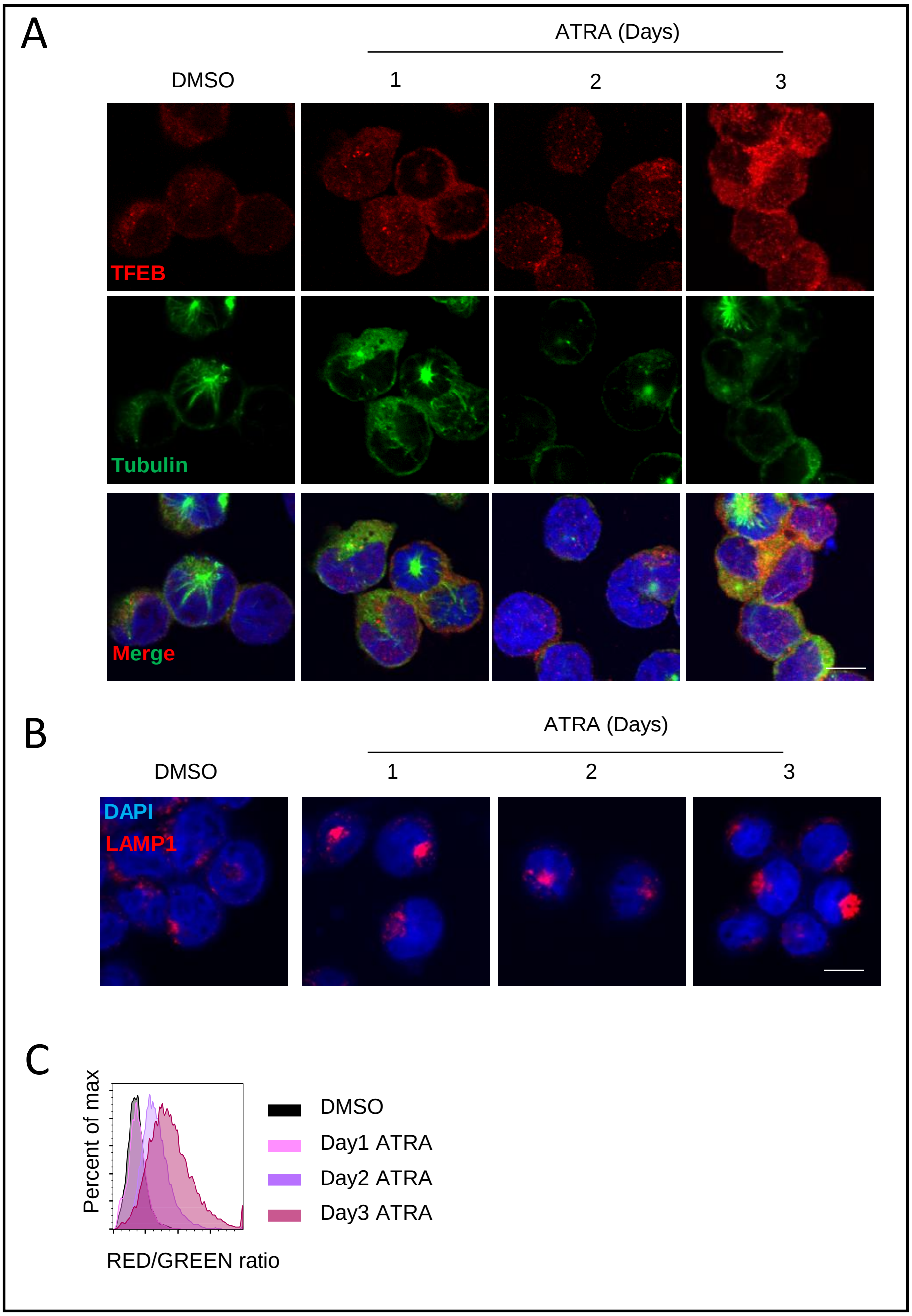

### Supplementary fig 6

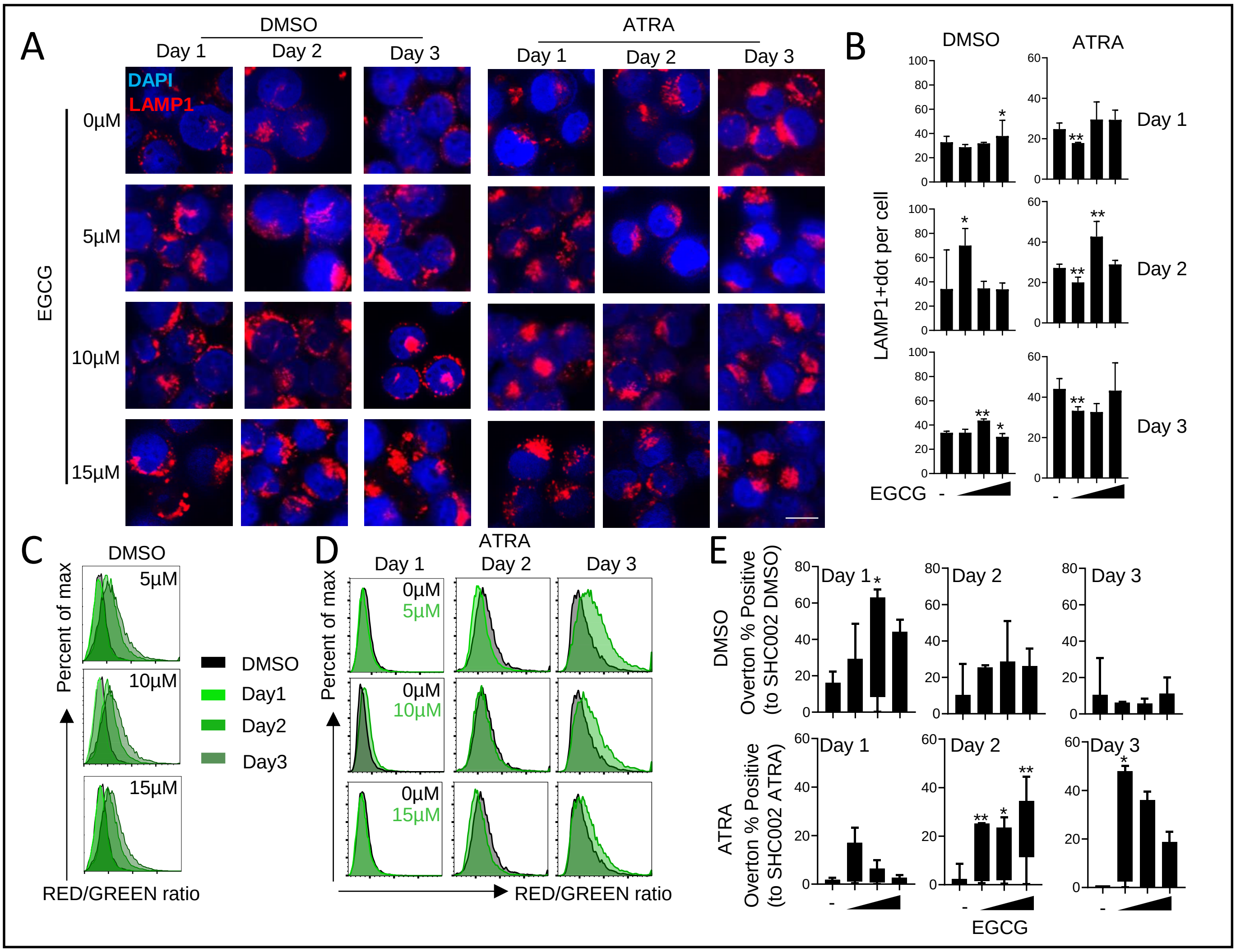
